# AlphaFold predictions are valuable hypotheses, and accelerate but do not replace experimental structure determination

**DOI:** 10.1101/2022.11.21.517405

**Authors:** Thomas C. Terwilliger, Dorothee Liebschner, Tristan I. Croll, Christopher J. Williams, Airlie J. McCoy, Billy K. Poon, Pavel V. Afonine, Robert D. Oeffner, Jane S. Richardson, Randy J. Read, Paul D. Adams

**Affiliations:** New Mexico Consortium, Los Alamos, NM 87544, USA; Los Alamos National Laboratory, Los Alamos, NM 87545, USA; Molecular Biophysics & Integrated Bioimaging Division, Lawrence Berkeley National Laboratory, Berkeley, CA 94720, USA; Department of Haematology, Cambridge Institute for Medical Research, University of Cambridge, Hills Road, Cambridge CB2 0XY, United Kingdom; Department of Biochemistry, Duke University, Durham, North Carolina, 27710; Department of Bioengineering, University of California, Berkeley, Berkeley, CA 94720, USA

## Abstract

AI-based methods such as AlphaFold have revolutionized structural biology, often making it possible to predict protein structures with high accuracy. The accuracies of these predictions vary, however, and they do not include ligands, covalent modifications or other environmental factors. Here we focus on very-high-confidence parts of AlphaFold predictions, evaluating how well they can be expected to describe the structure of a protein in a particular environment. We compare predictions with experimental crystallographic maps of the same proteins for 102 crystal structures. In many cases, those parts of AlphaFold predictions that were predicted with very high confidence matched experimental maps remarkably closely. In other cases, these predictions differed from experimental maps on a global scale through distortion and domain orientation, and on a local scale in backbone and side-chain conformation. Overall, C_α_ atoms in very-high-confidence parts of AlphaFold predictions differed from corresponding crystal structures by a median of 0.6 Å, and about 10% of these differed by more than 2 Å, each about twice the values found for pairs of crystal structures containing the same components but determined in different space groups. We suggest considering AlphaFold predictions as exceptionally useful hypotheses. We further suggest that it is important to consider the confidence in prediction when interpreting AlphaFold predictions and to carry out experimental structure determination to verify structural details, particularly those that involve interactions not included in the prediction.

---

Protein structure predictions using AlphaFold^1^, RoseTTAFold^2^, and related methods^3^ are far more accurate than previous generations of prediction algorithms^4^, bringing much closer to reality the biological understanding that could be derived from knowing the three-dimensional structures of all macromolecules^1,2,5-9^. AlphaFold predictions have already been made available for 200 million individual protein sequences to further drug discovery, protein engineering and understand biology^10^. A question that immediately arises is to what extent these predictions can substitute for experimental structure determinations^11,12^.

Both experimentally-determined protein structures and predicted models have important limitations^11,13,14^. Proteins are flexible and dynamic, and their distributions of conformations depend on temperature, solution conditions, and binding of ligands or other proteins (including crystal contacts in the case of crystallography)^15^. A model of a high-resolution crystal structure can accurately represent the dominant conformation(s) present in a crystal in a particular environment^11^, but the structure may differ under another set of conditions^14^. AI-based models can in many cases be very accurate, however they do not yet take into account the presence of ligands, covalent modifications or environmental factors, and take protein-protein interactions and multiple conformations into account in a limited way^1,2,16,17^.

The accuracy of a prediction is typically assessed by how closely it matches a structure in the Protein Data Bank^18^ (PDB) with the same sequence, and there are many ways to make such a comparison^4^. Using comparisons that focus on local accuracy, predictions obtained with AlphaFold have been assessed as having “atomic accuracy”^19^, as having accuracies competitive with “the best experimental results”^4^ and being of comparable quality as an experimental crystal structure^7^. It has been argued that AlphaFold predictions might be more accurate than estimated by comparison with models in the PDB, or even more accurate than the deposited models, because the deposited models are poorly defined in some places^4^. This reasoning notes that side-chain positions and loops are sometimes not clear in crystallographic electron density maps^20^, and in such cases a difference between an AlphaFold prediction and a deposited model would not indicate an error in the prediction. On the other hand, analyses carried out by the DeepMind team and others show that AlphaFold predictions vary substantially in their global and local agreement with deposited models and also in their coverage at the highest levels of confidence^1,11,21^, with only 36% of residues in the human proteome^22^ and 73% of residues in *E. coli* modeled with very high confidence^23^. Of course many of the proteins in the human proteome that have low-confidence AlphaFold predictions are likely to be regions that are intrinsically disordered^24,25^ and those would therefore also often not be revealed by experimental methods.

Here we address the accuracies of AlphaFold predictions by assessing how well they agree with experimental data^26^. We put these results into context by examining how closely one crystal structure in the PDB can typically be reproduced by another crystal structure containing the same components, but crystallized in a different space group (resulting in different crystal contacts).

## Comparing AlphaFold predictions with crystallographic electron density maps

We used a set of crystallographic electron density maps determined without reference to deposited models as standards for evaluation of AlphaFold predictions. The density maps were obtained^27^ using iterated AlphaFold prediction and model rebuilding with X-ray crystallographic data deposited in the PDB. For the present work we selected a high-quality subset of 102 models and maps from this analysis consisting of those that had free R values of 0.30 or better. The density maps in our analysis do not have any bias towards deposited models, as no information from deposited structures was used to compute these maps. Therefore, if features of a prediction are incompatible with the density maps and different from the deposited model, they are likely to be incorrect representations of the actual molecule in the crystal.

AlphaFold predictions are produced with residue-specific confidence metrics (pLDDT) which are estimates of the local accuracy of the prediction^1^. Residues with pLDDT values of greater than 90 are considered to be predicted with very high confidence and those with values of 70 or greater have moderate-to-high-confidence.

Figure 1 compares AlphaFold predictions, experimental density maps, and corresponding deposited models (predictions were superimposed on the deposited models). All the residues shown in Fig. 1 were predicted with very high confidence (pLDDT > 90) and the density maps range in resolution from 1.1 Å to 1.6 Å.

**Figure 1.**
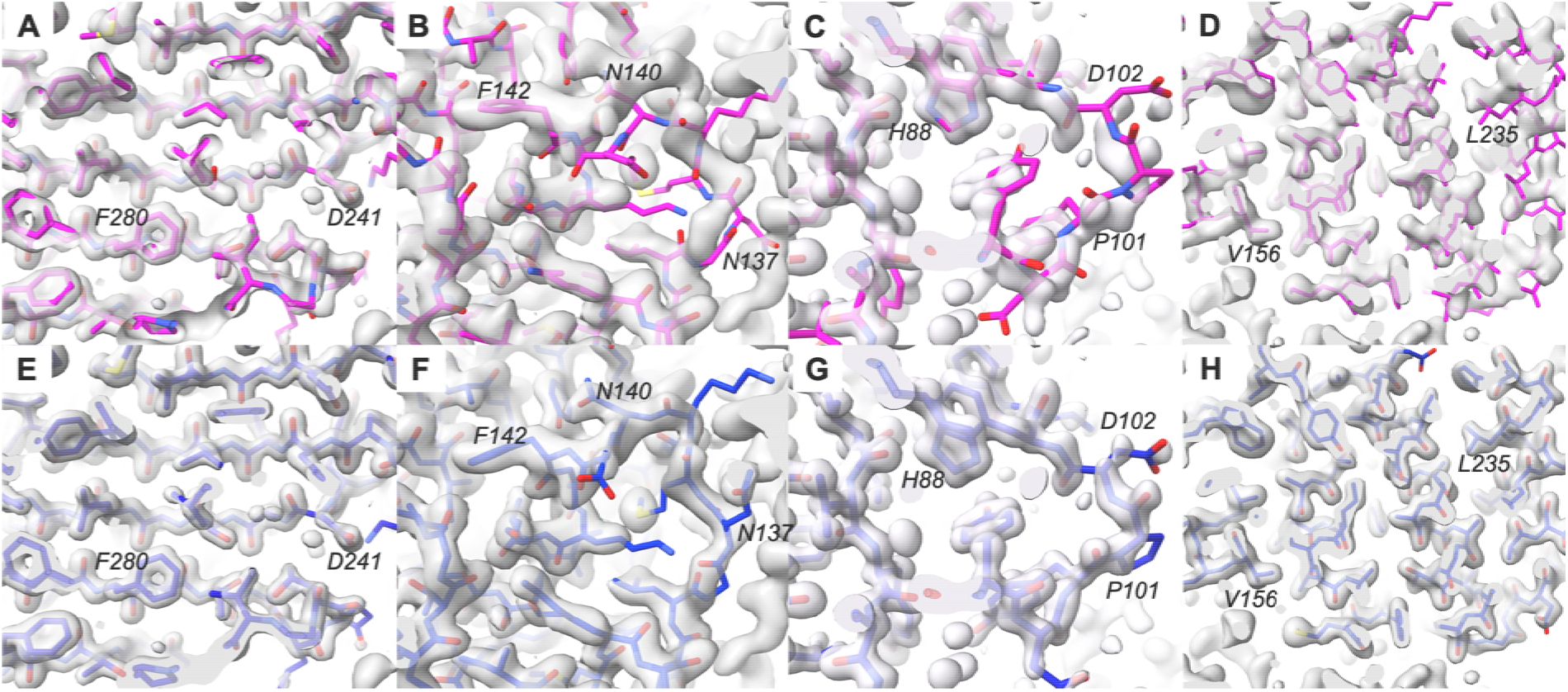
Comparison of details of AlphaFold predictions with density maps. AlphaFold predictions are shown in magenta with selected residues labeled; deposited models are shown in blue. Experimental electron density maps were taken from our previous work^28^ and are contoured at 1.9 σ (**A**), (**E**), 1.1 σ (**B**), (**F**), 1.5 σ (**C**), (**G**), and 1.2 σ (**D**), (**H**). Model coloring is bright for parts of the models outside the density contours and dimmed for parts that are inside the contours. (A) and (E): PDB entry 7waa showing a region with high-accuracy prediction. (B) and (F): PDB entry 7s5L showing a region with incorrect prediction. (C) and (G): PDB entry 7t26 showing a prediction that does not match the density map, but where the density map is not fully clear. (D) and (H): PDB entry 7naz, showing a prediction that is distorted relative to the density map.

Figure 1A shows an example of an AlphaFold prediction that superimposes closely on the corresponding density map (PDB entry 7waa^29^). For comparison, Fig. 1E shows the deposited model along with the same density map. The overall map-model correlation for the superimposed AlphaFold prediction is 0.72 and the rms C_α_ difference from the deposited model is 0.5 Å.

Figure 1B shows a prediction for PDB entry 7s5L^30^ which contained high-confidence regions that did not match the density map. The main chain corresponding to residues N137 through F142 match the density map poorly. In contrast, the deposited model matches the map very closely (Fig. 1F). The overall map-model correlation for the superimposed prediction is 0.44, much lower than that for the 7waa prediction shown in Fig. 1A, and the rms C_α_ difference from the deposited model is 2.1 Å.

Figure 1C shows an example of a prediction that does not match the density map but that might still represent a plausible conformation of the molecule. The prediction for PDB entry 7t26^31^ does not superimpose on the density near P101 and D102, while the deposited model does (Fig. 1G). The density map is less clear in this region than in other parts of the map. A break in main-chain density at D102 suggests that the chain adopts multiple conformations in this region. It is possible that the conformation in the AlphaFold prediction could be one of these alternative conformations, though not a dominant one as it does not appear in the density map.

Figure 1D illustrates a case where the AlphaFold prediction is distorted relative to the density map (PDB entry 7naz). Residues in the vicinity of V156 match the density closely (Fig. 1D), while residues near L235 are shifted relative to the map. For comparison, the deposited model matches the map closely throughout the region shown (Fig. 1H).

Figure 2A (open bars) shows the overall compatibility of 102 AlphaFold predictions with their corresponding density maps, as measured by map-model correlation. The mean map-model correlation for AlphaFold predictions (open bars) after superimposing them on corresponding deposited models was 0.56, substantially lower than the mean map-model correlation of deposited models to the same maps of 0.86 (hatched bars).

**Figure 2.**
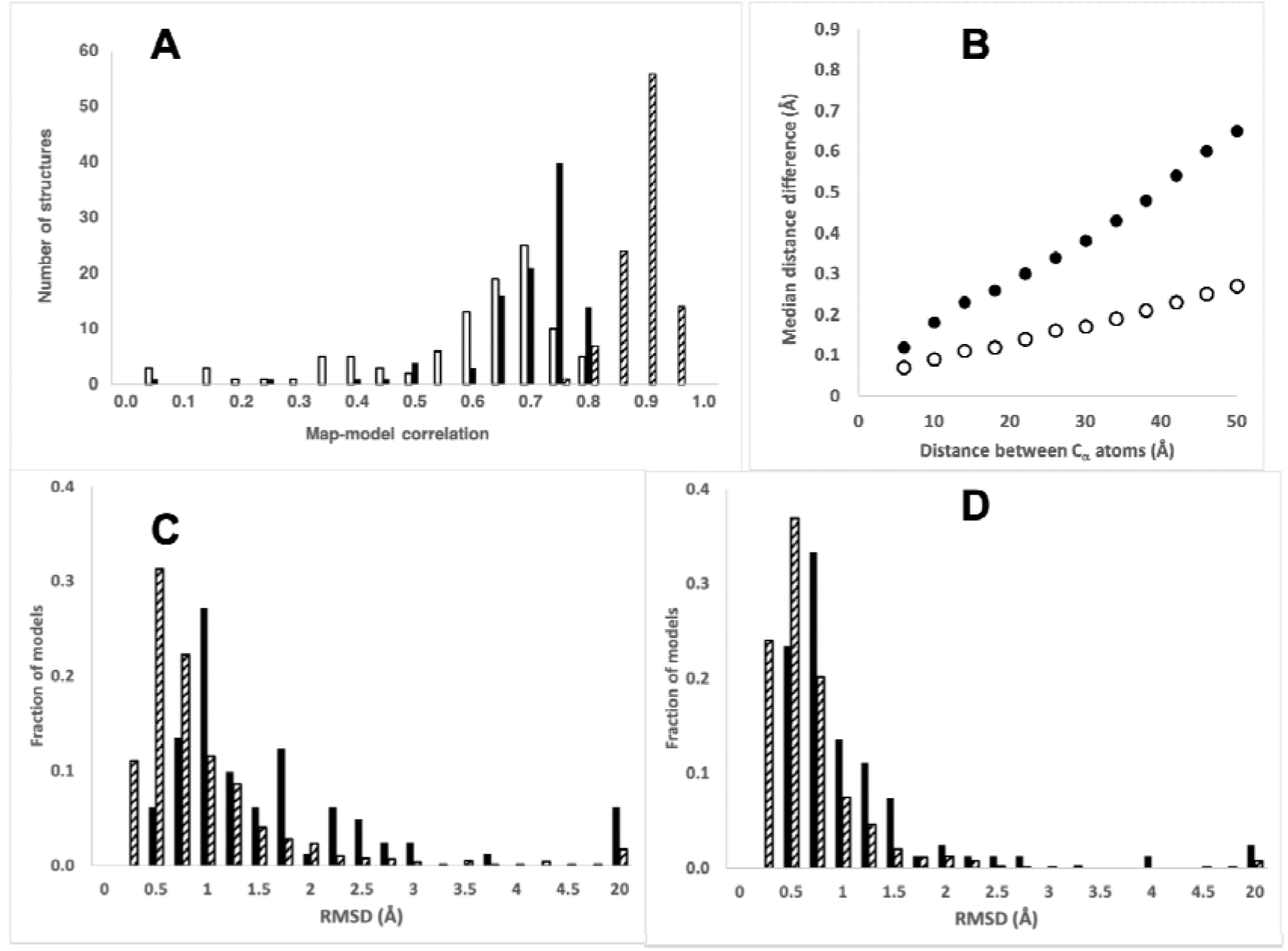
Overall comparison of AlphaFold predictions with density maps and deposited models. (**A**): Map-model correlation between 102 AlphaFold predictions (open bars), morphed AlphaFold predictions (solid bars), or corresponding deposited models (hatched bars) and experimental density maps. (**B**): Filled circles, median differences between distances in 102 AlphaFold predictions and those in corresponding deposited models, binned by the C_α_ - C_α_ distances (bin width of 4 Å). Open circles, as filled circles, but comparing matched pairs of structures from the PDB in which the components are the same but the crystal form is different. (**C**): RMSD between AlphaFold predictions and deposited models (solid bars) and between pairs of matching PDB entries with the same composition (hatched bars). The category at the far right on the abscissa labelled “20” includes all values greater than 5 Å. (**D**): As in C except after morphing models to match.

### Distortion and domain movement in AlphaFold predictions

Figure 1D illustrated that an AlphaFold prediction can be somewhat distorted relative to the actual structure. To determine whether this occurs for many AlphaFold predictions, we “morphed” each AlphaFold prediction to make it more similar to the deposited model (see Materials and Methods). This process reduces differences between predictions and deposited models that arise from either distortion or alternate locations of domains within chains. After morphing each predicted model, the predictions agree more closely with the electron density maps (Fig. 2A, solid bars, mean map correlation of 0.67 vs 0.56 before morphing), but still much less closely than the deposited models (Fig. 2A, hatched bars, mean map correlation of 0.86).

If two models are related by a long-range distortion or alternate locations of domains, inter-atomic distances that are short will be similar in the two models, while those that are long will differ. We quantified this relationship by comparing inter-atomic distances in predicted models with matching distances in deposited models and examining the median differences as a function of distance. Fig. 2B shows that this median inter-atomic distance deviation between deposited models and moderate-to-high-confidence parts of AlphaFold predictions (pLDDT above 70) is about 0.1 Å for atom pairs that are close (between 4 Å and 8 Å apart) and increases to 0.7 Å for distant atom pairs (48 Å – 52 Å), indicating a typical distortion of about 0.5-1 Å over this range of distances. As a reference, we analyzed 926 pairs of high-resolution structures in the PDB that had identical sequences but were obtained in different crystallographic space groups (so that crystal contacts influencing conformation would differ). Fig. 2B shows that atom pairs in these matching structures had distances that differed by a rms of 0.1 Å for nearby residues and 0.4 Å for distant ones, about half the values found for AlphaFold predictions.

As a third method of assessing distortion and differences in domain relationships in AlphaFold predictions, we compared them with the corresponding models from the PDB, calculating the rmsd of C_α_ atoms both before and after applying the distortion field described above. For this analysis we used all 215 structures analyzed in our previous work^28^. Fig. 2C shows the distribution of C_α_ rmsd values for the AlphaFold predictions; the median rmsd is 1.0 Å. After applying the distortion field, the median rmsd is reduced to 0.4 Å (Fig. 2D, the median rsmd distortion applied was 0.6 Å). For matching pairs of structures in the PDB crystallized in different space groups, the median C_α_ rmsd was only 0.6 Å, and this could be reduced to 0.4 Å by applying a distortion field (median rms distortion applied of 0.2 Å). Overall, the C_α_ coordinates in AlphaFold predictions are considerably more different from PDB entries than deposits of high-resolution structures of the same molecule in different space groups are from each other (median rmsd of 1.0 Å vs 0.6 Å), and a substantial part of this difference consists of long-range distortion.

### Comparing AlphaFold side-chain predictions with experimental density maps

As illustrated in Fig. 1, AlphaFold predictions often contain at least some regions that are similar to deposited structures, but even in these regions many details often differ. We used the 102 electron density maps described above along with deposited models to evaluate side-chain conformations (the locations of atoms in side chains relative to the atoms in the main chain that they are connected to), an important local feature of a structural model. In order to analyze the local side-chain structure and remove confounding effects from domain shifts or distortions, we grafted the side chain from each residue in an AlphaFold prediction onto the corresponding main chain atoms residue of the deposited model. This yielded a composite model with the main-chain coordinates of the deposited models and side-chain conformations corresponding to the AlphaFold predictions.

Figure 3A shows a local portion of PDB entry 7vgm, and Fig. 3B shows the AlphaFold prediction superimposed on the deposited model. Fig 3C shows the same region with the grafted side chain and the composite model. The positions of several of the side chains in the AlphaFold model (e.g., R32, D62, E530, E533, R494) are different from those in the deposited model. Fig. 3D shows the deposited model for 7vgm along with the density map obtained for PDB entry 7vgm, and Fig. 3E shows the AlphaFold model superimposed on the same density map. Even though the density map was obtained with the AlphaFold prediction and without reference to the deposited model, all the side chains in the deposited model match the map closely. In contrast, side chains in the AlphaFold prediction that were different from those in the deposited model do not match the density map, both before (Fig. 3E) and after (Fig. 3F) grafting, indicating that these side-chain conformations are likely to be incorrect.

**Figure 3.**
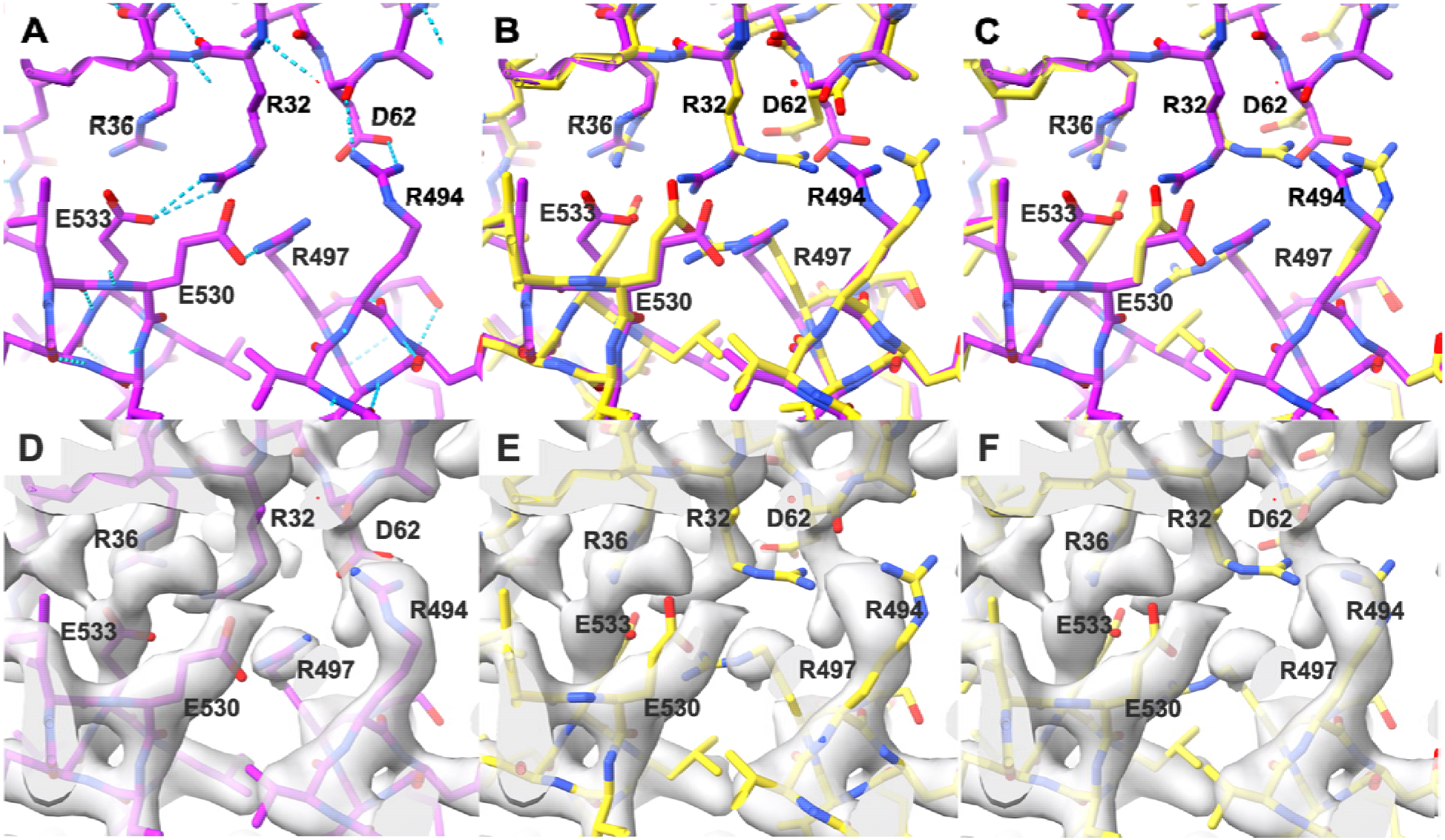
Comparison of AlphaFold side-chain predictions with density map for PDB entry 7vgm. (**A**): PDB entry 7vgm showing hydrogen bonding network. (**B**): AlphaFold prediction (yellow) superimposed on deposited model for PDB entry 7vgm (magenta). (**C**): As in B, except the AlphaFold side chains (yellow) are grafted on to the backbone for PDB entry 7vgm (main-chain atoms for each model are used to superimpose the side chains). (**D**): Deposited model as in A superimposed on experimental density map (2.3 Å resolution). (**E**): AlphaFold prediction as in B superimposed on density map. (**F**): grafted AlphaFold model superimposed on density map.

We carried out this side-chain grafting procedure for 102 AlphaFold predictions and the corresponding deposited models. For each pair of side chains, we examined the agreement between atomic positions in that side chain and the corresponding optimized density map. We identified pairs in which the AlphaFold side-chain prediction differed substantially from the deposited model (rmsd of side-chain atoms > 1.5 Å). Then based on estimates of the uncertainty of density values in each map and of the number of independent points sampled by side-chain atomic positions in that map, we identified AlphaFold side-chain predictions that differed from the deposited model and were highly unlikely (p < 0.01) to be as compatible with the density map as the deposited model. We considered these AlphaFold side-chain predictions to be incompatible with the experimental data.

Overall, we found that 20% of the side chains in moderate-to-high confidence residues of AlphaFold predictions and not involved in crystal contacts had different conformations than in the corresponding deposited model (at least 1.5 Å rmsd), and one third of these (7% overall) were clearly incompatible with the experimental data. As the number of clearly-incompatible residues identified by our method is a lower estimate, we expect that the actual level of disagreement between AlphaFold predictions and conformations of the molecules in the crystals is somewhere between the 7% that are clearly incompatible with the data and the 20% that differ from the deposited models.

To put the fraction of side-chain positions in AlphaFold predictions that are incompatible with the experimental data into perspective, we carried out a similar analysis, but using the set of high-resolution structures from the PDB containing the same components but crystallized in a different space group. For these tests we used experimentally-based density maps (2mFo-DFc maps^32^ calculated using one model from each pair. Here, only 6% of the side chains differed by 1.5 Å rmsd, and only 2% were in conformations that were experimentally incompatible with the corresponding conformations from the other set. Therefore, at a detailed level as well as an overall level, the differences between AlphaFold predictions and these crystal structures are substantially greater than for pairs of crystal structures determined in different space groups.

We then analyzed whether the 7% of residues in AlphaFold predictions that were incompatible with experimental data included residues with functional significance. We extracted all the residues that were explicitly mentioned in the 49 publications describing the 102 analyzed structures, yielding a total of 733 named residues. Of these, 53 (7%) were among the residues we identified as being incompatible with experimental data, the same percentage that we found for all residues. For example, residues R32, D62, R497 and E533 in Fig. 3 are all in this group of functional residues that are incompatible with experimental data.

As functionally significant residues are constrained by evolution, it might have been expected that the evolutionary covariation that forms a central element of AlphaFold prediction^19^ would be stronger than average. On the other hand, these same residues are more conserved than average^33^, possibly balancing that effect. In our small sample, we do not see a substantial effect either way, rather we find that side chains for residues in AlphaFold predictions with functional significance are about as likely to be incompatible with experimental data as other side chains.

### Using confidence (pLDDT) to estimate errors in AlphaFold predictions

As AlphaFold predictions can differ substantially from corresponding experimental models, straightforward methods to estimate coordinate uncertainties of these predictions would be useful. As a first step, we superimposed AlphaFold predictions on corresponding deposited models and determined the distance between the C_α_ atoms in the predicted and deposited models, as well as the confidence (pLDDT) for the predicted C_α_ atom.

Figure 4A shows the distribution of prediction errors for various ranges of the confidence measure. For comparison, the dashed line in Fig. 4A shows the distribution of differences between matching C_α_ atoms in pairs of structures containing the same components but crystallized in different space groups. The median prediction error for high-confidence (pLDDT > 90) residues was 0.6 Å, while for residues with pLDDT between 80 and 90 it was 1.1 Å, and for those between 70 and 80 it was 1.5 Å (Table I). By comparison, matching C_α_ atoms in pairs of structures in different space groups differed by a median of 0.3 Å. Fig. 4B shows that morphing one member of each pair as described above reduces the differences over all confidence ranges, but differences between matching pairs of structures in the PDB are reduced similarly.

**Figure 4.**
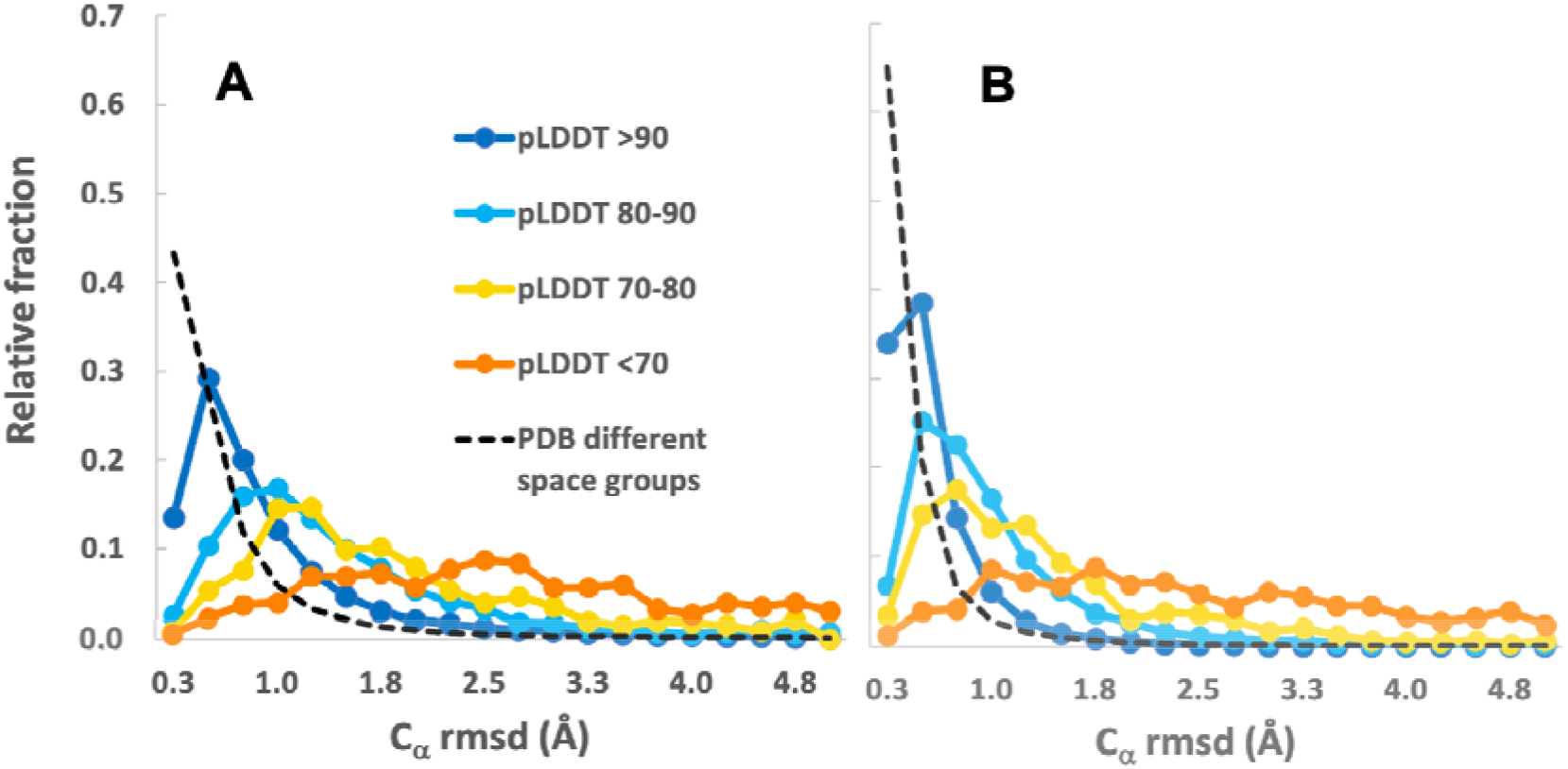
Distribution of prediction errors for ranges of AlphaFold prediction confidence. Dark blue dots and line, pLDDT > 90, light blue, between 80 and 90, yellow, between 70 and 80, orange, less than 70. Ordinate is the fraction of cases in the ranges of rmsd indicated on the abscissa. Dashed line shows similar comparison for matching pairs of PDB deposits with different space groups. (**A**): errors estimated for structures as is. (**B**): Errors estimated after morphing.

**Table I.** Median prediction error and percentage with prediction error over 2 Å by AlphaFold confidence.

| AlphaFold confidence (pLDDT) | Median prediction error (Å) | Percentage with error over 2 Å |
| --- | --- | --- |
| >90 | 0.6 | 10 |
| 80 - 90 | 1.1 | 22 |
| 70 - 80 | 1.5 | 33 |
| <70 | 3.5 | 77 |

The significance of the median coordinate errors found above depends on what the coordinates are going to be used for^13,14^. If coordinates are intended for use in comparing distantly-related structures to infer evolutionary and structural relationships, where typical differences among structures may be large (for example, 2 – 3 Å), median coordinate errors of 1.1 Å may have little effect on the analysis. On the other hand, the same coordinate errors might substantially affect an analysis involving docking of a ligand to identify specific protein-ligand interactions.

We note the distributions in Fig. 4 do not resemble the Maxwell-Boltzmann distribution expected for random 3-dimensional Gaussian errors (there is an excess kurtosis of over 200 for errors in prediction vs an expected value of 0.1). The distributions have a small fraction of values that are very large (long tails in the distributions), so describing uncertainties in terms of rms errors may not ordinarily be effective. Instead, it may be more useful to note the median errors described above as a measure of typical errors, and to also take into account the percentage of instances where the error is very large (i.e., completely wrong). The definition of very large errors will depend on the situation, but often atomic positions that deviate by more than 2 or 3 Å are of limited value.

For the structures analyzed here, about 10% of C_α_ atoms with pLDDT over 90 are found to be in error by over 2 Å, along with 22% of those with pLDDT between 80 and 90, 33% of those between 70 and 80, and 77% of those with pLDDT under 70 (Table I). For comparison, just 5% of C_α_ atoms in the matched pairs of structures in the PDB crystallized in different space groups we analyzed differ by over 2 Å.

The extent of agreement between AlphaFold predictions and experimental data found here is consistent with results of the uncertainty quantification carried out by DeepMind during the development of AlphaFold^25^. That analysis showed that estimated model accuracy (pLDDT) was an unbiased predictor of actual model accuracy (LDDT), and that the correlation between pLDDT estimates and actual LDDT was about 0.76. The uncertainty quantification further estimated that 7% (for pLDDT > 90) to 30% (for 70 < pLDDT < 90) of side chains have a χ_1_ angle deviation of at least 40°. Such a deviation typically leads to an rmsd of side-chain atoms of over 1.5 Å. In our analysis, the average pLDDT was 94, with 12% of residues having a pLDDT between 70 and 90. Therefore, the errors estimated in AlphaFold development are generally consistent with our observation that between 7% and 20% of side chains with pLDDT of 70 or above are incompatible with experimental data.

## Conclusions

While AlphaFold predictions are often astonishingly accurate (e.g., Fig 1A), we find that many parts of AlphaFold predictions are incompatible with experimental data from corresponding crystal structures. In particular, our results show that AlphaFold predictions are not better representations of the contents of a crystal than the models deposited in the PDB, as the deposited models agree much more closely with experimental data where the predicted and deposited models differ. Our results also show that even very high confidence AlphaFold predictions differ from corresponding models deposited in the PDB by about twice as much as pairs of high-resolution structures in the PDB that were crystallized in different space groups, indicating that AlphaFold predictions are in error by more than the amount that might be expected due to flexibility. We note that as AlphaFold prediction does not take into account the presence of ligands, ions, covalent modifications, or environmental conditions, it cannot be expected to correctly represent the many details of protein structures that depend on these factors.

A confidence metric (pLDDT) is produced for each AlphaFold prediction. This confidence metric was examined in detail by the DeepMind team and was shown to be unbiased (equally likely to be too low or too high) and to have a good correlation to the LDDT metric that it estimates (Pearson’s correlation of 0.76)^1^. This confidence metric can therefore be a very useful residue-specific indicator of the accuracy of a prediction. For the structures examined here, the parts of AlphaFold predictions that had very high confidence (pLDDT > 90, 86% of residues in the analysis) were generally quite accurate (median C_α_ coordinate difference from deposited model of 0.6 Å). It is important to note, however, that about 10% of residues predicted with very high confidence differed from the deposited model by over 2 Å (Table I).

Despite their limitations, AlphaFold predictions are already changing the way that hypotheses about protein structures are generated and tested^1,2,5,6^. Indeed, even though not all parts of AlphaFold predictions are accurate, they provide plausible hypotheses that can suggest mechanisms of action and allow designing experiments with specific expected outcomes. Using these predictions as starting hypotheses can also greatly accelerate the process of experimental structure determination^27,34,35^. AlphaFold predictions often have very good stereochemical characteristics, making them excellent hypotheses for local structural features. For example, for the 102 structures analyzed here, the mean percentage of residues with “favored” Ramachandran configurations was 98%, greater than that of the corresponding deposited models (97%), and the mean percentage of side-chain conformations classified as outliers was just 0.2%, compared with 1.5% for deposited models^28^. Such AlphaFold predictions with highly plausible geometry could be used in later stages of experimental structure determination as potential conformations for segments of structure that are not fully clear in experimental density maps.

All these capabilities are very likely just the beginning of an age of increasingly broad use of AI methods in structural biology^12^. AI approaches will surely be extended from proteins to include nucleic acids, ligands, covalent modifications, environmental conditions, interactions among all these entities, and multiple structural states. The accuracies of these predictions and of the uncertainties associated with them are very likely to improve continuously as additional factors are included and as databases of sequence and structural information expand. The resulting predictions will be increasingly useful structural hypotheses that will form a solid foundation for experimental and theoretical analyses of biological systems.

## Funding

Lawrence Berkeley National Laboratory grant DE-AC02-05CH11231 (PDA)

National Institutes of Health grant GM063210 (PDA, JSR, RJR, TCT)

Wellcome Trust grant 209407/Z/17/Z (RJR)

Phenix Industrial Consortium (PDA)

## Author contributions

Conceptualization: TCT, PDA, RJR, JSR

Methodology: TCT, PDA, RJR, JSR

Investigation: TCT, AJM, BKP, PVA, TIC, CJW, DL, RDO

Visualization: TCT

Funding acquisition: TCT, PDA, RJR, JSR

Project administration: TCT, PDA, RJR, JSR

Supervision: TCT, PDA, RJR, and JSR

Writing – original draft: TCT

Writing – review & editing: TCT, PDA, RJR, JSR, AJM, BKP, PVA, TIC, CJW, DL, RDO

## Data and materials availability

Input data for deposited models were taken from the Protein Data Bank. The 102 accession codes used were: 7e0m, 7fhr, 7v6p, 7Ljh, 7p3a, 7v38, 7v3b, 7o9p, 7rLz, 7qdv, 7ewj, 7rw4, 7waa, 7kdx, 7fiu, 7n3v, 7ptb, 7dtr, 7aoj, 7rc2, 7tcr, 7wja, 7vnx, 7x8v, 7raw, 7rpy, 7aov, 7tb5, 7t8L, 7vwk, 7ne9, 7nqd, 7s5L, 7wbk, 7x77, 7e3z, 7f0o, 7v1q, 7etx, 7ety, 7ecd, 7dxn, 7eyj, 7e4d, 7wsj, 7fi3, 7wnn, 7vgm, 7eio, 7v9n, 7tvc, 7Lbk, 7e6v, 7b3n, 7bLL, 7djj, 7dms, 7dqx, 7drh, 7dri, 7e1d, 7e85, 7edc, 7ejg, 7es4, 7esi, 7eus, 7ew8, 7exx, 7f2a, 7fjg, 7kzh, 7Lsv, 7mku, 7naz, 7ncy, 7nxg, 7o51, 7o5y, 7oc3, 7oom, 7oq6, 7qs4, 7rm7, 7t7j, 7tbs, 7tem, 7tfq, 7tj1, 7tL5, 7tmu, 7tog, 7toj, 7trv, 7trw, 7tt9, 7twc, 7tzp, 7unn, 7w3s, 7wdq, 8cuk. All models are downloadable from the PDB with links such as: https://files.rcsb.org/download/7tzp.pdb or (for larger models that are not available in this format) https://files.rcsb.org/download/7tzp.cif. We used the Phenix tool *fetch_pdb* to download models and crystallographic data for each structure. Predicted models, rebuilt models, and density-modified map coefficients are available at: https://phenix-online.org/phenix_data/terwilliger/alphafold_crystallography_2022/ along with a spreadsheet that contains all the raw data and analyses described in our previous work^28^ and described here. The directory terwilliger/alphafold_crystallography_2022/contains a README file describing the contents of the site, the spreadsheet, and a data/directory with one compressed archive for each structure containing models and crystallographic data files. This directory also contains a compressed archive (*alphafold_crystallography*.*tgz*) containing all the data and all the scripts used to create the spreadsheet.

## Code Availability

All code for the Phenix version of the AlphaFold2 Colab is freely available on GitHub at https://github.com/phenix-project/Colabs. All code for Phenix is available at phenix-online.org.

## Supplementary Materials

### Materials and Methods

#### Experimental data, models, AlphaFold predictions, and density maps

We used the results of our automated structure redeterminations^28^ for crystallographic PDB deposits in this work. The structures in that study were chosen based on the method of structure solution (single-wavelength anomalous diffraction, SAD), used as a proxy for relatively challenging structure determinations. The anomalous data were not used in our structure redeterminations, i.e., the Bijvoet pairs were averaged. All the unique, protein-containing structures in a 6-month period (Dec. 2021-May 2022) were analyzed (215 structures). Structures were determined with molecular replacement using trimmed AlphaFold predictions^36^ as search models, followed by iterative model rebuilding and AlphaFold prediction^27^. In this work we use the initial AlphaFold predictions (made without templates) and the final density-modified electron density maps^37^ from those analyses. Except as noted, in this work we used only structures yielding a free R value of 0.30 or lower (102 structures) to ensure that the density-modified electron density maps used as a reference were of high quality.

#### Model morphing with a distortion field

We used a morphing procedure based on a smoothed distortion field^38^ to modify one model to make it globally more similar to another model, while retaining local differences. In this procedure any point in space has an associated shift vector, the shift that is to be applied to any atom located at that point in space. This association of a vector to each point in space amounts to a shift or distortion field. To create a smoothly-varying distortion field relating a pair of structures, we first create an exact distortion field that maps one structure onto the other, then this field is smoothed.

First, the two structures are superimposed. Then a set of positions in space and corresponding shift vectors is created, with the positions in space ***y***_*i*_ corresponding to C_α_ atom coordinates in one structure, and the shift vectors ***v***_*i*_ corresponding to the differences between matching C_α_ atoms in the two structures. At this point, each of these positions in space has the property that if the associated shift vector is added, it will match the corresponding C_α_ atom coordinate in the other structure. This exact distortion field is defined only at the C_α_ atom coordinates of the first structure.

Then we create a smoothed distortion field ***v(x)*** that is defined at any point in space ***x*** by averaging all the shift values in the exact distortion field, weighting individual shifts ***v***_*i*_ with a weight ***w***_*i*_ based on the distances between their positions in space ***y***_*i*_ and that point ***x***,

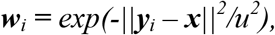

where the scaling factor *u* determines the distance over which smoothing occurs, typically set to 15 Å.

#### Analytical procedures

Map-model correlations for predicted models were calculated after superposition on the corresponding deposited models.

For structures with more than one chain, only the first chain was included for each structure in comparisons.

#### Side-chain grafting

The grafting procedure was carried out using the *model_building*.*graft_side_chains* method in Phenix. This function identifies matching residues in two models, then uses the coordinates of atoms in the main chain for a residue in one model to position the main-chain and side-chain atoms in a matched residue from another. We excluded residues with low confidence (pLDDT < 70, 2% of the total residues), and residues that participate in crystal contacts (any atom in the residue within 6 Å of any atom in a symmetry-related molecule, 23% of all residues).

#### Choice of examples of AlphaFold predictions with varying relationships to density maps (Fig. 1)

The goal of Fig. 1 is to illustrate four situations that occurred among the AlphaFold predictions that we examined. We noticed four distinct situations (prediction agrees exceptionally well with density map, prediction does not match density map, prediction does not match density map but might be correct, prediction is distorted relative to the map). We then chose one example of each type that was as clear as possible and that contained only very-high-confidence parts of these predictions to display.

#### Evaluation of compatibility of side-chain positions with density maps

We identified side-chain conformations in AlphaFold predictions that were incompatible with corresponding electron density maps as cases where the predicted side-chain conformation matched the density map much more poorly than the deposited model and differed substantially from that found in the corresponding deposited model. To focus on the side-chain conformation separately from the overall location and orientation of each residue, we used the side-chain grafting procedure described above to orient the main chain of each residue from an AlphaFold prediction to match the main chain of the corresponding residue in the deposited model. We considered side chains to differ substantially if the rmsd of side-chain atoms beyond the C_β_ atom was greater than 1.5 Å.

We then identified incompatible AlphaFold side-chain conformations as those that were highly unlikely (p < 0.01) to be as compatible with the density map as the deposited model. This probability was estimated from the uncertainty of density values in each map and the number of independent points sampled by side-chain atomic positions in that map. To obtain the uncertainty of density values, we calculated the rms difference between *Fobs* and *Fcalc* maps obtained from the *phenix*.*refine*^*39*^ software using the deposited model and crystallographic data to calculate the maps. To estimate the number of independent points sampled by side-chain atomic positions for a particular side chain, we counted the number of side-chain atoms that could be selected where each atom is separated from all others by at least the resolution of the data.

As an example of this procedure, for the 7vgm example shown in Fig. 3, the mean electron density map value at atoms in the side chain of residue R32 in 7vgm was 2.8 and the mean density for the side chain from the AlphaFold prediction was 0.1 (the map is normalized to have a mean of zero and rms of 1). These side chains differed by an rmsd of 3.9 Å and the 6 side-chain atoms corresponded to approximately 4 unique positions in the map (4 positions that are each separated from the others by the resolution of the map). The map, adjusted to have a mean of zero and rms of 1, had an estimated uncertainty of 0.8 (based on agreement between the calculated and observed structure factor amplitudes), leading to a probability of p < 10^−10^ that the AlphaFold prediction is actually in better agreement with the map than the deposited model.

## Supplementary text

### Control experiments and limitations

Our analysis of side-chain conformations is based on the premise that the backbone conformation of the deposited model is largely correct. However, it is possible that the backbone is systematically distorted at residues with incorrect rotamers, as the main chain atom positions might compensate for errors in the side chain. We checked for this scenario by refitting the side chains for all 102 structures, and applying a “backrub” correction to the main chain to correct for these distortions if necessary^40^. A repeat of our analysis, skipping the 4% of side chains where a backrub correction was applied (C_β_ shift^40^ of more than 0.2 Å), yielded very similar results, with 18% of residues differing in side-chain orientation and again 7% overall clearly incompatible with experimental data.

We also checked for the possibility that backbone conformations might differ in the two models for some residues, making the grafting procedure inappropriate. We repeated our analysis, removing all residues where the Ramachandran angles differed in the two structures by more than 30° (10% of all residues). Once again, the results were similar, with 17% of residues differing in side-chain orientation and 7% overall clearly incompatible with experimental data.

Our test set (102 for most analyses, 215 for some) is a small fraction of those in the entire PDB, so it could be useful to analyze a larger, more representative set. Most of the residues in our analysis had very high confidence, with 86% having pLDDT values above 90, 10% from 80 to 90, 2% from 70 to 80 and 2% under 70. In contrast, in the AlphaFold prediction of the human proteome^25^, only 36% of residues had pLDDT values above 90, and 42% were under 70. The small fraction of residues with predictions under 80 may lead to some uncertainty in the error estimates for moderate and low-confidence predictions in Table I. The median rmsd between AlphaFold predictions and deposited models in the PDB in our analysis (1.0 Å, see Fig. 2C in main text) was considerably lower than that obtained in a large-scale analysis of recent structures by DeepMind^1^ (2.3 Å for all C_α_ atoms, 1.5 Å excluding the largest 5% of differences), perhaps due to the high confidence in prediction in our sample.

As we wanted to estimate the accuracy of the 200 million predictions made with the standard version, we did not remove predictions that might be better-predicted with a multimer version of AlphaFold^16^. For example, PDB entry 7e1d is a domain-swapped dimer^41^ that was predicted by AlphaFold to be a compact chain.

In some instances, domain-swapping or other incorrect connections between domains resulted in very large differences between predictions and deposited models. Therefore, we attempted to reduce the effect of these outlier structures by quoting median values where possible.

We used a local installation of AlphaFold for our predictions and did not use templates from the PDB in prediction, which could reduce the accuracy of the predicted models. Based on a comparison of our AlphaFold predictions and those in the AlphaFold database^10^, which included templates in prediction, this effect is likely to be small, however. We identified 81 models in the AlphaFold database that corresponded to the first chains in one of our 102 analyses. The median C_α_ atom rmsd between our initial predicted models^28^ and the corresponding chain in the AlphaFold database was just 0.54 Å. The predictions from the AlphaFold database had a median rmsd of 1.15 Å compared to deposited models; our predictions without templates also had an rmsd of 1.15 Å.

## References

1 Jumper, J. et al. Highly accurate protein structure prediction with AlphaFold. Nature 596, 583–589, (2021).

2 Baek, M. et al. Accurate prediction of protein structures and interactions using a three-track neural network. Science 373, 871–876, (2021).

3 Lin, Z. et al. Evolutionary-scale prediction of atomic level protein structure with a language model. bioRxiv, 2022.2007.2020.500902, (2022).

4 Kryshtafovych, A., Schwede, T., Topf, M., Fidelis, K. & Moult, J. Critical assessment of methods of protein structure prediction (CASP)—Round XIV. Proteins: Structure, Function, and Bioinformatics 89, 1607–1617, (2021).

5 Callaway, E. ‘The entire protein universe’: AI predicts shape of nearly every known protein. Nature 608, 15–16, (2022).

6 Thornton, J. M., Laskowski, R. A. & Borkakoti, N. AlphaFold heralds a data-driven revolution in biology and medicine. Nature Medicine 27, 1666–1669, (2021).

7 van Breugel, M., Rosa e Silva, I. & Andreeva, A. Structural validation and assessment of AlphaFold2 predictions for centrosomal and centriolar proteins and their complexes. Communications Biology 5, 312, (2022).

8 Subramaniam, S. & Kleywegt, G. J. A paradigm shift in structural biology. Nature Methods 19, 20–23, (2022).

9 Ourmazd, A., Moffat, K. & Lattman, E. E. Structural biology is solved — now what? Nature Methods 19, 24–26, (2022).

10 Hassabis, D. AlphaFold reveals the structure of the protein universe, https://www.deepmind.com/blog/alphafold-reveals-the-structure-of-the-protein-universe

11 Shao, C., Bittrich, S., Wang, S. & Burley, S. K. Assessing PDB macromolecular crystal structure confidence at the individual amino acid residue level. Structure, (2022).

12 Goulet, A. & Cambillau, C. Present Impact of AlphaFold2 Revolution on Structural Biology, and an Illustration With the Structure Prediction of the Bacteriophage J-1 Host Adhesion Device. Frontiers in Molecular Biosciences 9, (2022).

13 Moore, P. B., Hendrickson, W. A., Henderson, R. & Brunger, A. T. The protein-folding problem: Not yet solved. Science 375, 507, (2022).

14 Acharya, K. R. & Lloyd, M. D. The advantages and limitations of protein crystal structures. Trends in Pharmacological Sciences 26, 10–14, (2005).

15 Fraser, J. S. et al. Accessing protein conformational ensembles using room-temperature X-ray crystallography. Proceedings of the National Academy of Sciences 108, 16247–16252, (2011).

16 Evans, R. et al. Protein complex prediction with AlphaFold-Multimer. bioRxiv, 2021.2010.2004.463034, (2022).

17 Stein, R. A. & McHaourab, H. S. SPEACH_AF: Sampling protein ensembles and conformational heterogeneity with Alphafold2. PLoS Comput Biol 18, e1010483, (2022).

18 consortium, w. Protein Data Bank: the single global archive for 3D macromolecular structure data. Nucleic Acids Res 47, D520–D528, (2018).

19 Jumper, J. & Hassabis, D. Protein structure predictions to atomic accuracy with AlphaFold. Nature Methods 19, 11–12, (2022).

20 van Beusekom, B., Joosten, K., Hekkelman, M. L., Joosten, R. P. & Perrakis, A. Homology-based loop modeling yields more complete crystallographic protein structures. IUCrJ 5, 585–594, (2018).

21 Hryc, C. F. & Baker, M. L. AlphaFold2 and CryoEM: Revisiting CryoEM modeling in near-atomic resolution density maps. iScience 25, 104496, (2022).

22 Porta-Pardo, E., Ruiz-Serra, V., Valentini, S. & Valencia, A. The structural coverage of the human proteome before and after AlphaFold. PLOS Computational Biology 18, e1009818, (2022).

23 Akdel, M. et al. A structural biology community assessment of AlphaFold2 applications. Nat Struct Mol Biol 29, 1056–1067, (2022).

24 Dunker, A. K. et al. Intrinsically disordered protein. Journal of Molecular Graphics and Modelling 19, 26–59, (2001).

25 Tunyasuvunakool, K. et al. Highly accurate protein structure prediction for the human proteome. Nature 596, 590–596, (2021).

26 Flower, T. G. & Hurley, J. H. Crystallographic molecular replacement using an in silico-generated search model of SARS-CoV-2 ORF8. Protein Science 30, 728–734, (2021).

27 Terwilliger, T. C. et al. Improved AlphaFold modeling with implicit experimental information. Nature Methods, (2022).

28 Terwilliger, T. C. et al. Accelerating crystal structure determination with iterative AlphaFold prediction. bioRxiv, 2022.2011.2018.517112, (2022).

29 Zhang, Q. et al. Re-sensitization of mcr carrying multidrug resistant bacteria to colistin by silver. Proc Natl Acad Sci U S A 119, e2119417119, (2022).

30 Burkhardt, I., de Rond, T., Chen, P. Y.-T. & Moore, B. S. Ancient plant-like terpene biosynthesis in corals. Nature Chemical Biology 18, 664–669, (2022).

31 Hobbs, S. J. et al. Phage anti-CBASS and anti-Pycsar nucleases subvert bacterial immunity. Nature 605, 522–526, (2022).

32 Read, R. Improved Fourier coefficients for maps using phases from partial structures with errors. Acta Crystallographica Section A 42, 140–149, (1986).

33 Bartlett, G. J., Porter, C. T., Borkakoti, N. & Thornton, J. M. Analysis of Catalytic Residues in Enzyme Active Sites. Journal of Molecular Biology 324, 105–121, (2002).

34 McCoy, A. J., Sammito, M. D. & Read, R. J. Implications of AlphaFold2 for crystallographic phasing by molecular replacement. Acta Crystallographica Section D 78, 1–13, (2022).

35 Barbarin-Bocahu, I. & Graille, M. The X-ray crystallography phase problem solved thanks to AlphaFold and RoseTTAFold models: a case-study report. Acta Crystallogr D Struct Biol 78, 517–531, (2022).

36 Oeffner, R. D. et al. Putting AlphaFold models to work with phenix.process_predicted_model and ISOLDE. Acta Crystallographica Section D 78, (2022).

37 Terwilliger, T. Maximum-likelihood density modification. Acta Crystallographica Section D 56, 965–972, (2000).

38 Cowtan, K., Metcalfe, S. & Bond, P. Shift-field refinement of macromolecular atomic models. Acta Crystallographica Section D 76, 1192–1200, (2020).

39 Afonine, P. V. et al. Towards automated crystallographic structure refinement with phenix.refine. Acta Crystallographica Section D 68, 352–367, (2012).

40 Davis, I. W., Arendall, W. B., Richardson, D. C. & Richardson, J. S. The Backrub Motion: How Protein Backbone Shrugs When a Sidechain Dances. Structure 14, 265–274, (2006).

41 Bennett, M. J., Choe, S. & Eisenberg, D. Domain swapping: entangling alliances between proteins. Proc Natl Acad Sci U S A 91, 3127–3131, (1994).

